# A transient ischemic environment induces reversible compaction of chromatin

**DOI:** 10.1101/025221

**Authors:** Ina Kirmes, Aleksander Szczurek, Kirti Prakash, Iryna Charapitsa, Christina Heiser, Michael Musheev, Florian Schock, Karolina Fornalczyk, Dongyu Ma, Udo Birk, Christoph Cremer, George Reid

## Abstract

The environmental effects of ischemia on chromatin nanostructure were evaluated using single molecule localisation microscopy (SMLM) of DNA binding dyes. Short-term oxygen and nutrient deprivation (OND) of the cardiomyocyte cell-line HL-1induces a previously undescribed chromatin architecture, consisting of large, chromatin sparse voids interspersed between DNA-dense hollow helicoid structures of the order of 40 to 700 nm in dimension. OND induced chromatin compaction is reversible, and upon restitution of normoxia and nutrients, chromatin transiently adopts a significantly more open structure than in untreated cells. We show that this compacted state of chromatin reduces transcription, while the open chromatin structure following recovery has a higher transcriptional rate than in untreated cells. Digestion of chromatin with DNAseI and DNA binding dye loading assays confirm that OND induces compaction of chromatin and a general redistribution of chromatin to the nuclear periphery. Mechanistically, chromatin compaction is associated with a depletion of intracellular ATP and a redistribution of the cellular polyamine pool into the nucleus. Additionally, Fluorescence Recovery After Photobleaching (FRAP) shows that core histones are not displaced from compacted chromatin and that the mobility of linker histone H1 is considerably reduced by OND treatment, to an extent that far exceeds the difference in histone H1 mobility between heterochromatin and euchromatin. These studies exemplify the dynamic capacity of chromatin architecture to physically respond to environmental conditions, directly link cellular energy status to chromatin compaction and provide insight into the effect ischemia has on the nuclear architecture of cells.

## Background

Cellular oxygen insufficiency, hypoxia, occurs in physiological and developmental processes and in disease, such as tumour growth, stroke and cardiac infarction. Hypoxia in pathological situations often results from ischemia and is associated with a reduced availability of glucose. Cells detect, accommodate and adapt to hypoxic and nutritional stress, and upon restoration of normoxia and nutrients, through immediate transcriptional, translational and metabolic responses. The major transcriptional mediator in hypoxia is the alpha/beta-heterodimeric hypoxia-inducible factor (HIF), which is present only when intracellular oxygen is low [1]. HIF activates the expression of genes involved in oxygen transport, glucose uptake, glycolysis and angiogenesis [2, 3]. Additionally, ischemia induced hypoglycemia results in stimulation of AMP-activated protein kinase (AMPK), a stress sensor which induces catabolic pathways and down-regulates anabolic processes such as fatty acid oxidation, glucose uptake and glycolysis upon cellular energy insufficiency [4, 5]. Moderate periods of hypoxia and nutrient deprivation provoke a predominant global repression of transcription [6, 7], although activation of hypoxia and/or hypoglycemic responsive genes does occur within this general transcriptionally repressive environment [8]. The transcriptional competence of DNA in eukaryotic cells is determined by its organization in chromatin. Chromatin structure is dynamically regulated at multiple levels, which include ATP-dependent chromatin remodelling [9], post-translation modification of chromatin [10] and the incorporation of histone variants [5].

The metabolic status of cells has a direct effect upon chromatin architecture, as many histone modifying enzymes utilize essential metabolites, such as ATP, NAD^+^, acetyl-coenzyme A (acetyl-CoA), S-adenosylmethionine (SAM) or oxygen either as cofactors or as substrates [11]. In particular, histone acetylation is dependent upon the action of ATP-citrate lyase [12], which converts mitochondrially derived citrate into cytoplasmically available acetyl-CoA. Additionally, molecular oxygen is required as a substrate by the Jumonji C (JmjC) class of dioxygenases to achieve histone demethylation. Consequently, hypoxia can limit the activity of a subset of JmjC histone demethylases, resulting in global increases in the methylation of histone H3K4, H3K9, H3K27 and H3K36 and in chromatin condensation [13]. For example, moderate hypoxia is reported to induce an overall decrease of H3K9 acetylation [14], with ischemia shown to decrease histone H4K16 acetylation in neural cells [15].

Nuclear architecture is dynamic and represents the structural and topological product of epigenetic regulation (for reviews see [16–18]). Chromosomes occupy distinct territories (CTs) in the cell nucleus [18–20], which harbour chromatin domains (CDs) with a size range in the order of 100 kbp to 1 Mbp [21-23]. In turn, CDs form chromatin domain clusters (CDCs) with a compact core and a less compact periphery, called the perichromatin region (PR) [24-26]. Histone marks associated with transcriptionally silent chromatin are enriched in the interior of CDCs, whereas marks typical for transcriptionally competent chromatin, and for chromatin associated with transcribing RNA polymerase II are enriched in the PR, where nascent RNA is synthesized [25-28]. CDCs, in turn, form a higher order chromatin network, which is attached to the nuclear envelope and permeates the nuclear interior. This chromatin network is co-aligned with a second network, called the interchromatin compartment (IC), which starts at nuclear pores [25, 29]. It pervades the nuclear space between CDCs and is enriched in proteins involved in genomic output. Previous work has demonstrated that chromatin architecture physically responds to environmental conditions, with condensation occurring in response to hyperosmotic conditions [30] and in response to oxidative stress provoked by the fungal metabolite chaetocin [31]. Depletion of ATP in HeLa cells results in chromatin compaction as evaluated by Fluorescence Lifetime Imaging Microscopy - Förster Resonance Energy Transfer (FLIM-FRET) [32]. Reflecting this, stress-induced and developmentally induced changes of gene expression correspond with major changes of nuclear organization [33]. As ischemia provokes major changes in transcriptional output, in the post-translational modification of histones and a reduction in intracellular ATP levels, it can be anticipated that OND may lead to significant changes in nuclear architecture.

While various biochemical approaches exist that evaluate the compaction state of chromatin, for example, chromatin capture technology [34], they do not report the underlying three dimensional nuclear structure. Recent advances in super-resolution optical microscopy provide structural discrimination comparable to that of electron microscopy [35]. Currently, single molecule localization microscopy (SMLM) has the highest spatial resolution of all optical microscopic methods used in cellular nanostructure analysis [36]. In the SMLM mode used here [37], most fluorophores are transferred to a metastable dark state, while a minor population of multiple emitting fluorophores remains that can be optically isolated and hence individually localised. In a typical determination, tens of thousands of frames are acquired over a period of several hours. Integrating the positions of fluorophores results in a joint localization map that can resolve spatial features of the order of 30 - 100 nm, in comparison to the approximately 250 nm limit of conventional optical methods [38, 39]. Direct imaging of DNA by means of localization microscopy is a prerequisite in determining chromatin structure, and has recently been accomplished for a range of DNA binding dyes [37, 40-42], (Żurek-Biesiada *et al.*, 2015, submitted).

We describe optically, at single molecule resolution, the effect of ischemic conditions on the nuclear architecture of immortalized cardiomyocytes. Exposure of HL-1 cells, a murine, adult cardiomyocyte cell-line [40] to moderate, acute hypoxia, (1% O_2_ for 1 hour), combined with nutrient starvation and glycolytic blockade, induces a condensed, hollow, whorl-like chromatin configuration with a concomitant reduction, around 30%, in the capacity of chromatin to associate with the DNA selective dye Vybrant DyeCycle Violet. Significantly, the occurrence of decondensed chromatin, as characterised by a diffuse distribution of DNA at the edge of chromatin territories, marked by the local presence of acetylated histones, is ablated and there is a marked relocation of condensed chromatin towards the nuclear periphery. Condensed chromatin exhibits an increased resistance to digestion with DNAseI, as compared to chromatin in untreated cells, and additionally, the mobility of linker histone H1, as estimated by FRAP, is significantly reduced by OND. Relaxation of nuclear architecture occurs within tens of minutes upon cessation of OND. Cytometric analysis of immunostained cells confirms and extends the findings made by single molecule localization studies. Mechanistically, chromatin compaction is associated with depletion of the intracellular pool of ATP, which results in the relocation of a substantial proportion of the cellular polyamine pool from the cytoplasm to the nucleus.

## Results

### Oxygen and nutrient deprivation of HL-1 cells induces chromatin compaction

We firstly assessed the response of chromatin to experimental conditions that mimic ischemia-reperfusion, through employing two-colour SMLM to characterise the response and recovery of nuclear architecture to transient OND in HL-1 cells, as evaluated by staining of DNA with DNA binding dyes and by immunodetection of H3K14ac, a histone mark associated with transcriptionally permissive chromatin. Fixed and permeablized cells were immunostained using AlexaFluor647 conjugated anti-H3K14ac and counter-stained with Vybrant DyeCycle Violet, a photoconvertible DNA binding dye which undergoes reversible photoswitching and that can be utilized for SMLM based on blinking, with individual fluorophores emitting up to 1500 photons per cycle (Żurek-Biesiada *et al.*, 2015, submitted). We typically generated localization maps, for at least nine nuclei per experimental condition, by integrating 30,000 observations, each of which captured photons emitted during a 50 ms exposure period. These observations localize individual fluorophores at sub-diffractional precision, with a theoretical lateral optical resolution of 67 nm and an experimentally determined structural resolution of 100 nm (Supplementary figure S10). As shown in figures 1a and inset b, untreated HL-1 cells show a typical DNA staining pattern, with rather intense staining occurring just inside the nuclear envelope and in discrete foci within the nucleus. There is a general diffuse staining of DNA within the nucleus, with small inter-nuclear compartments clearly visible between individual chromatin domains. H3K14ac occurs in a punctate distribution throughout the nucleus, with individual foci predominantly located at the edge of chromatin domains. This is in keeping with the topography found for the transcriptionally permissive H3K4me3 modification, in a range of mammalian cell types [25, 26]. The level of resolution and precision of localization obtained with our dual-colour SMLM technique cannot be attained by conventional microscopy (figure 1c).

**Figure 1.**
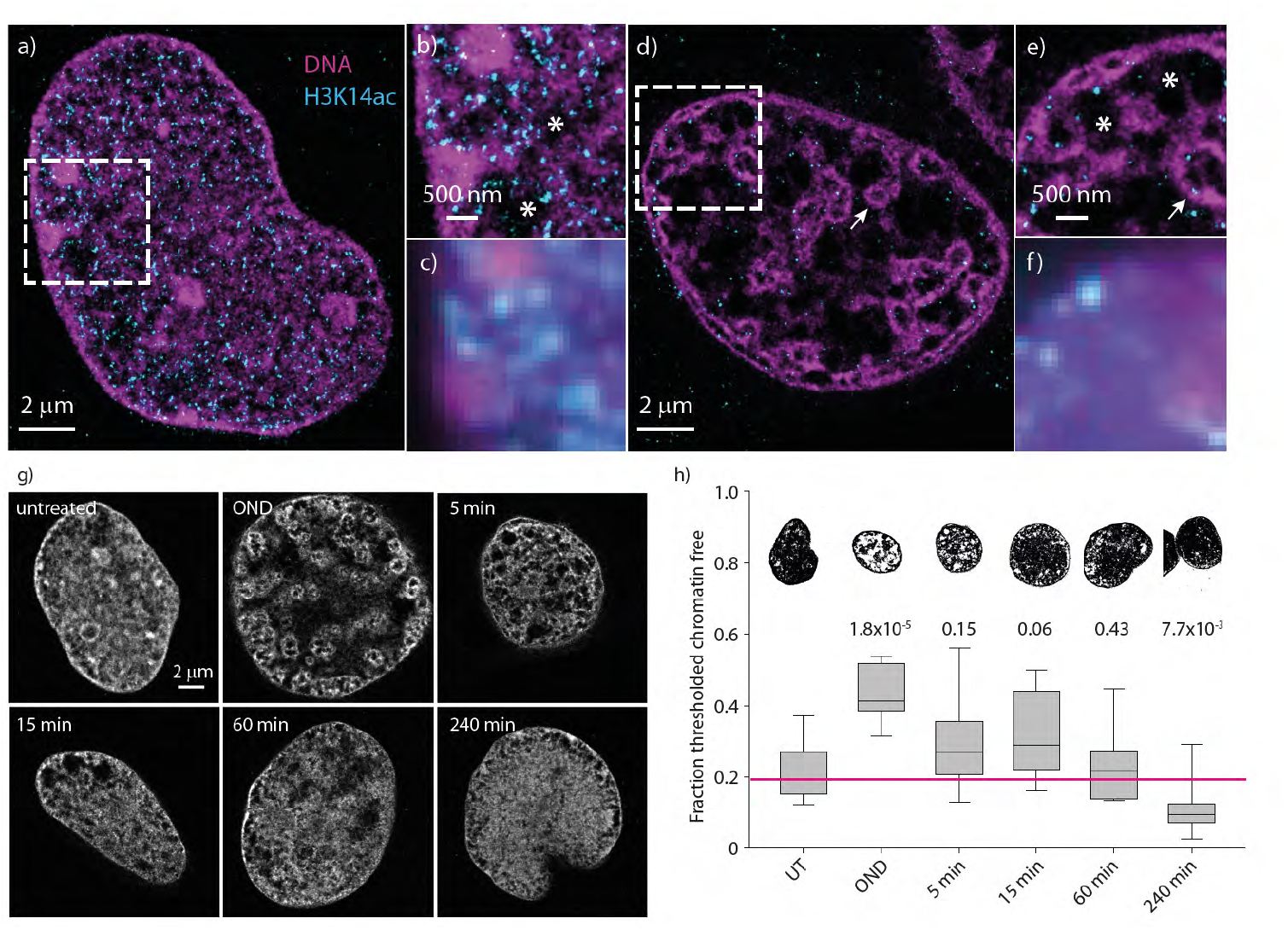
Oxygen and nutrient deprivation induces compaction of chromatin. HL-1 cells were fixed, permeabilized, immunostained with anti-acetylated histone H3K14 and then stained withVybrant DyeCycle Violet. Two colour SMLM was performed on untreated HL-1 cells (a and inset b) or on cells exposed for one hour of OND (d and inset e). For comparison, wide-field images of the inset regions are shown in c and in f. Chromatin voids are indicated by * with atolls marked by an arrow. Representative SMLM images of Vybrant Dyecycle Violet stained nuclei, either untreated, subject to one hour of OND or 5, 15, 60 and 240 minutes after release from OND are shown in panel g. A discriminatory threshold (pixel intensity ≤ 50) was applied to the experimental set of SMLM imaged nuclei (minimum of 9 cells were imaged), with boxplots and representative images describing the median and range of the proportion of the nucleus with chromatin shown in panel h, p-values compared to untreated are reported above the box-plots.

SMLM imaging of the nuclei of HL-1 cells exposed to one hour of OND demonstrates that an ischemic environment provokes a dramatic change in nuclear architecture, with condensed chromatin present at the sub-nuclear envelope, often as a closely spaced double arrangement of densely stained DNA, or as hollow intranuclear atolls (figure 1d and inset e). Moreover, the interchromosomal space consists of large, DNA sparse voids, with little of the diffuse DNA staining that is seen in untreated cells. A decrease in staining for H3K14ac occurs upon OND, with SMLM imaging again demonstrating that the remaining H3K14ac occurs largely at the edge of chromatin domains. In order to experimentally evaluate the effects of reperfusion following a transient ischemic period, we next evaluated the response of OND induced chromatin compaction to the restitution of normoxia and nutrients. SMLM of representative HL-1 cells either untreated, subject to one hour of OND or upon subsequent recovery from OND is shown in figure 1g. Following OND induced chromatin compaction, nuclear architecture relaxes and at 4 hours post-OND acquires a more open conformation than in untreated cells. To quantitatively evaluate this, we applied a discriminatory threshold to the experimental set of SMLM imaged cells, to delimit chromatin sparse nuclear regions. The distribution of nuclear areas that are chromatin-sparse is reported in figure 1h, with representative thresholded images shown above. OND induces an approximately two-fold increase in chromatin-free nuclear area. Sixty minutes of recovery from OND is sufficient for the majority of cells to restore chromatin architecture; however, a significant proportion of cells adopt a more open chromatin structure at 240 minutes. HL-1 cells recover fully from transient OND and continue to proliferate as well as untreated cells.

### Alternative staining and SMLM methodologies confirm that OND induces chromatin compaction

We then confirmed that OND induces extensive compaction of chromatin using an alternative nucleic acid binding dye, YOYO-1 [43] that also blinks under our experimental conditions, as previously reported [40] (figure 2, a-f) and with a click-chemistry approach chemically linking a fluorophore to 5-ethynyl-2′-deoxyuridine (EdU) [44] incorporated into DNA during cellular replication (figure 2, g-l). While both these approaches produce a reduction in the density of signals in comparison to Vybrant DyeCycle Violet, they clearly demonstrate that one hour of OND induces chromatin compaction in HL-1 cardiomyocytes.

**Figure 2.**
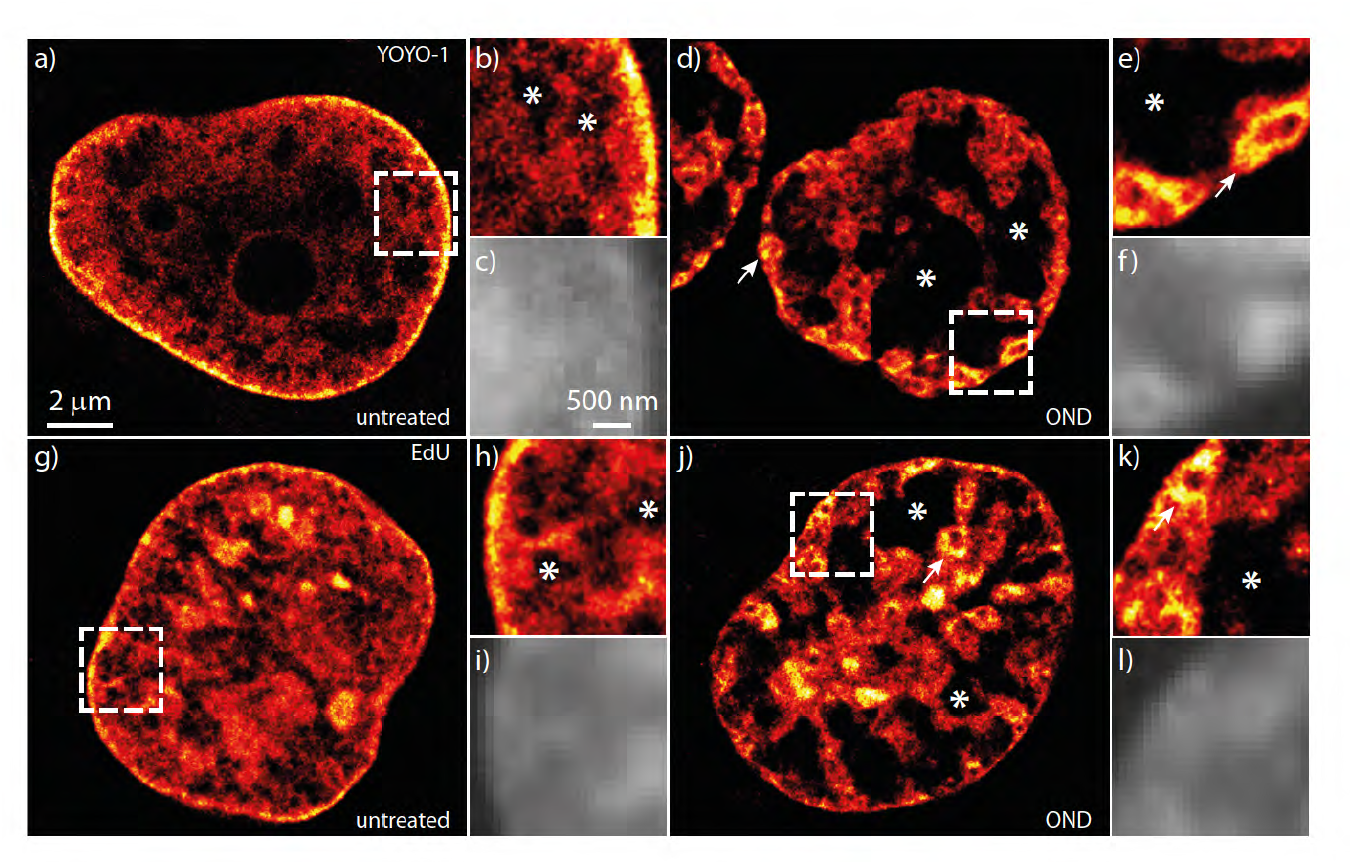
Alternative dyes and labelling methodologies confirm OND induced compaction of chromatin. HL-1 cells, either untreated (a-c) or exposed to one hour of OND (d-f) were fixed, permeabilized, stained with the DNA binding dye YOYO-1 and subject to SMLM (a, d and insets b, e). Alternatively, cells were labelled for 24 hours with 10 µM EdU, and then were either untreated (g-i) or subject to one hour of OND (j-l). Following fixation, EdU incorporated into DNA was coupled via click chemistry to AlexaFluor 488, as described [44] and nuclear DNA determined by SMLM (g, j and insets h, k) For comparison, wide-field images of the inset regions are shown in c, f, i and l. Chromatin voids are indicated by an *, with atolls marked by an arrow.

### A quantitative binning analysis describes the extent of chromatin compaction, the range of sizes of condensed structures and illustrates that chromatin adopts a more open structure on recovery from OND

SMLM defines the spatial localization of single fluorophores, permitting a quantitative assessment of chromatin condensation induced by OND. We initially evaluated the density of Vybrant DyeCycle Violet molecules detected by SMLM (figure 3a). Untreated cells have a median value of around 6×10^3^ dye localizations per µm^2^, which decreases by approximately 30% upon one hour of OND, and then recovers upon release from ischemia-mimetic conditions. Significantly, chromatin associates with approximately 30% more Vybrant DyeCycle Violet four hours after release from OND compared to untreated cells, again suggesting that chromatin adopts, at least transiently, a more open configuration upon recovery from ischemic conditions.

**Figure 3.**
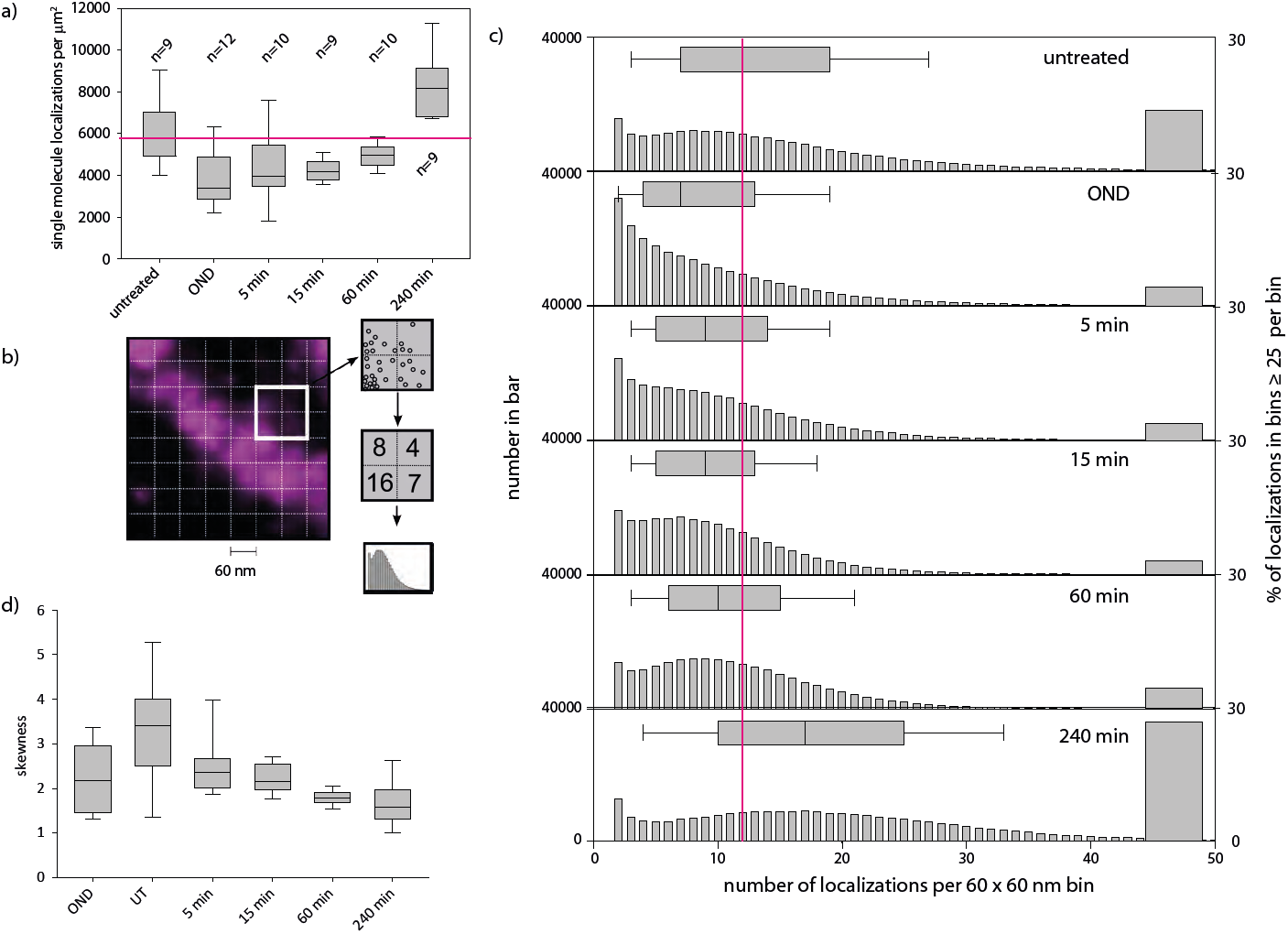
Quantification of chromatin compaction through binning. The influence of OND on the nuclear distribution and accessibility of chromatin was characterised by analysis of joint localization maps generated by SMLM. The median and range of densities of single molecule localizations, calculated over the entire nucleus and for a minimum of nine cells, for untreated, OND exposed and recovering cells, are presented as box plots in (a). A binning approach, outlined in (b), was then used to characterize the extent of chromatin compaction as cells transition from normal conditions through OND and in recovery from OND (c), with the median and range of the distribution shown above each histogram. The proportion of bins containing ≥ 25 localizations is presented as a bar on the right of each panel. As the distributions of histograms of binned data differ significantly between time points, the skewness (deviation from the mean) was calculated for all of the images and presented as a box plot (d).

We then used a binning approach to quantify the sub-nuclear distribution of chromatin, through counting individual SMLM locations of individual Vybrant Violet labelled DNA sites within a grid of squares (bins) overlaid upon the image of the nucleus (figure 3b). Reflecting the development of extensive regions within the nucleus that become chromatin sparse upon OND, the number of molecules present per bin becomes reduced in experimental ischemia, and this recovers upon restoration of normoxia and nutrients (figure 3c). A bin size of 60 × 60 nm was chosen to illustrate that this technique resolves structures at the tens of nm scale, with bins containing either 0 or 1 localization excluded from the presented results. Reflecting the increase in DNA dye binding seen four hours post OND, there is a corresponding increase at four hours in the proportion of bins containing a large number of localizations (figure 3c), indicating that recovery from OND induces chromatin to adopt, at least transiently, a more open conformation. In order to describe the spatial extent of changes of chromatin density induced by OND, we evaluated a range of bin sizes of between 10 to 500 nm for untreated and OND cells. We then evaluated the skewness, a measure of asymmetry around the mean, of the distributions throughout the experimental time-course (figure 3d) and found that the distribution of chromatin density become more skewed in the direction of high DNA density classes than in untreated cells (median skewness of ∼3.2 and ∼2.2 respectively). Notably, as the skewness parameter is positive in all experimental conditions, including untreated cells, it can be inferred that the majority of chromatin is located within a highly condensed state rather than in a diffuse conformation. Similar results were obtained for EdU-Alexa488 labelled chromatin (Supplementary Figure S11).

### Nearest neighbour analysis confirms and describes the extent of chromatin compaction

We further characterised chromatin condensation induced by OND, through determining the average distance of single molecule localizations to variable numbers of nearest neighbours within computationally tractable representative regions of interest (ROI). An example of selected ROI’s is shown in figure 4a. Three ROI’s from three independent nuclei were used to generate datasets for each experimental condition. We firstly evaluated the relationship between the average distance to nearest neighbours and the number of neighbours evaluated. The median distance and range of distribution rises with the number of neighbours used in the analysis (figures 4b and 4d). Reflecting chromatin compaction, the average distance to neighbours increases upon one hour of OND, which becomes resolved upon restoration of normoxia and energy source (figure 4c). These effects become more apparent when further adjacent neighbours are included in the analysis, at least up to 500 nearest neighbours (figure 4d).

**Figure 4.**
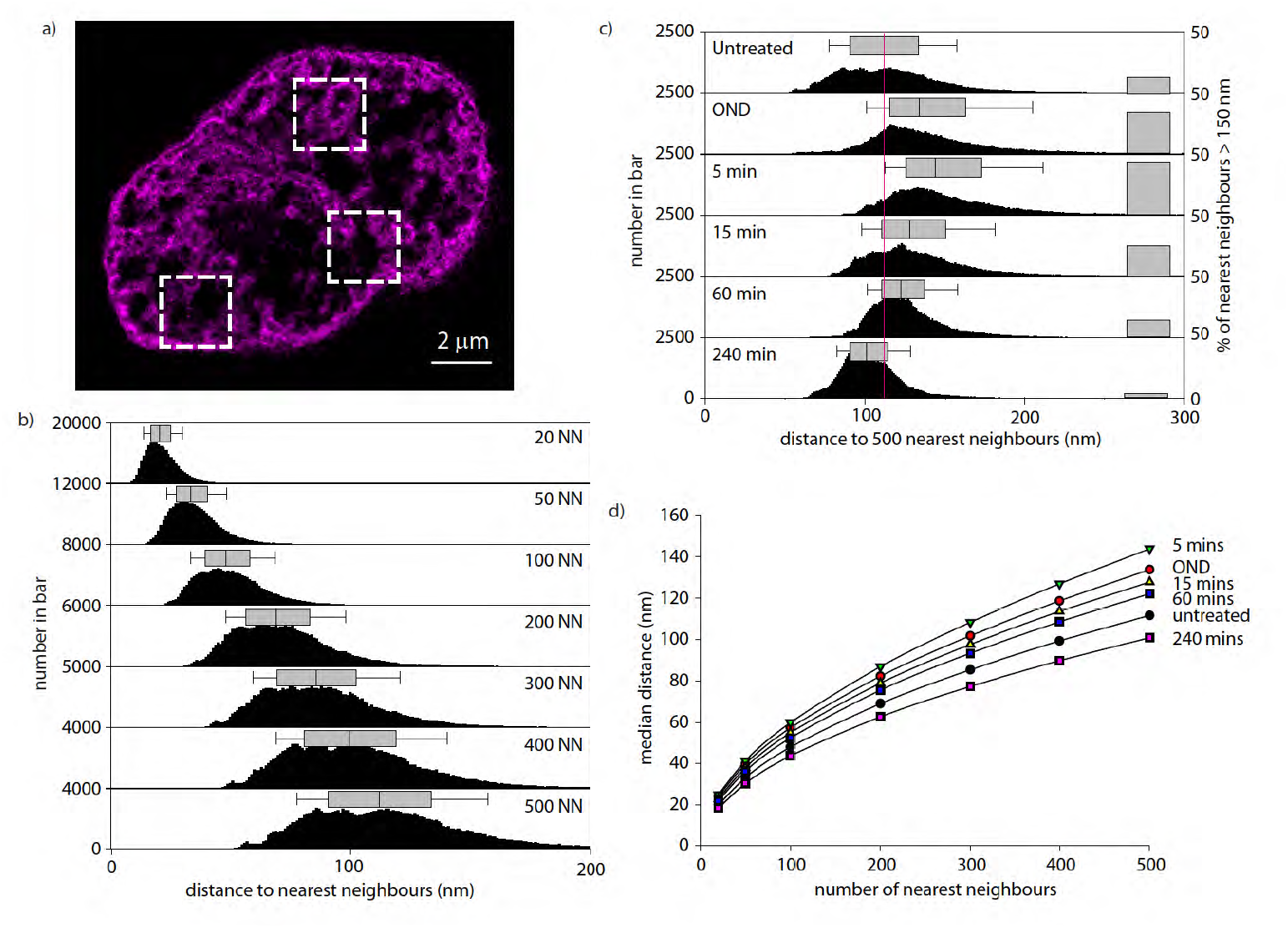
Nearest Neighbour characterization of OND induced chromatin compaction. Nearest neighbour analysis was used to describe the extent of chromatin compaction upon deprivation of oxygen and nutrients using three internal regions of interest (ROI), as illustrated for an HL-1 cell subject to one hour of OND (a). Results were generated for each experimental condition using three ROI’s per nucleus and three nuclei per determination. The effect of the number of nearest neighbours evaluated on the distance to the analysed position is shown in (b) as a histogram and as a box-plot showing the median and range of distribution of values for untreated cells. The extent of chromatin compaction as cells transition from normal conditions through OND and in recovery from OND is shown in (c), using the distance to 500 nearest neighbours, with the median and range of distribution shown above each histogram. The proportion of bins with a distance to 100 nearest neighbours ≥ 80 nm is shown as a bar on the right of each panel. The relationship between the number of nearest neighbours used in the analysis to the median distance to the set of nearest neighbours is shown for each experimental condition in (d).

### OND decreases the susceptibility of chromatin to DNAseI digestion

We next utilised a biochemical approach to confirm that OND treatment for one hour does indeed provoke compaction of chromatin. We estimated the access of a large molecular weight probe, DNAseI (30 kDa), to chromatin. DNA in fixed and permeabilised untreated or OND treated cells was preloaded for one hour with DRAQ5, a selective DNA interchelating dye [45], and then subject to digestion with DNAseI, with cellular fluorescence continuously measured on a confocal platform. Digestion of DNA provokes liberation of DRAQ5, with the rate of decrease in DRAQ5 fluorescence dependent upon on the extent of chromatin compaction. As shown in figure 5, untreated cells show a triphasic response to DNAseI treatment, with a highly accessible sub-fraction of chromatin, approximately 50% of the total, predominating the kinetics of the first 15 minutes of the timecourse. A more compact, but nevertheless digestible, fraction then defines the following 40 minutes of digestion, with a residual proportion of chromatin, some 10% of the total, predominantly resistant to DNAseI digestion. In contrast, OND cells show a biphasic response, with a compact but digestible fraction dominant for the first 60 minutes of digestion, followed by a fraction of chromatin, around 30% of the total that is relatively resistant to DNAseI digestion. OND cells do not exhibit a rapidly digested fraction of chromatin, as observed in untreated cells. These results confirm and extend our SMLM observations that oxygen and nutrient deprivation induces profound compaction of chromatin, particularly of loosely condensed chromatin.

**Figure 5.**
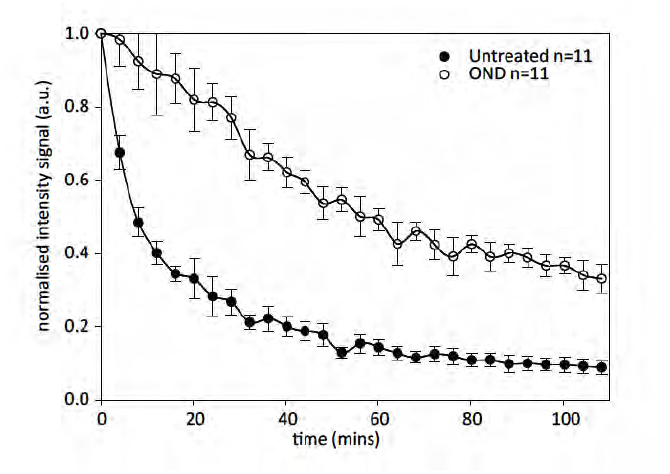
OND induces chromatin compaction as determined by resistance to digestion by DNAseI. HL-1 cells, either untreated or subject to one hour of OND, were fixed, permeabilised and stained with 5 µM DRAQ5 for 30 minutes. Cells were then digested with 5 U/ml DNAseI at 37oC with cellular fluorescence measured on a confocal microscope, with images generated every four minutes observing in total 11 cells for each experimental condition.

### OND reduces cellular ATP levels, inhibits transcription, redistributes polyamines to the nucleus and restricts access to histones

We then postulated that chromatin compaction induced by OND is consequent upon ATP depletion. Under normal conditions, divalent cations and polyamines associate with the polyphosphate group of ATP. However, if ATP levels are reduced, these may, by mass-action, relocate to the sugar-phosphate backbone of nucleic acid, thereby promote chromatin compaction through effecting charge shielding. OND reduces intracellular ATP levels by 90%, which recovers upon cessation of OND with kinetics similar to that of chromatin relaxation (figure 6a). Furthermore, OND promotes a global decrease of transcription by approximately 90%, as estimated by mass spectrometric determination of bromouridine incorporation into nascent RNA (figure 6b). We then described the distribution of the intracellular polyamine pool using immunocytochemistry. Anti-polyamine staining of untreated HL-1 cells results in a punctate, predominantly cytoplasmic distribution with a low level of intranuclear staining (figure 6c). This most likely reflects ATP rich mitochondria present in the cytoplasm of cardiomyocytes. In contrast, OND treatment for one hour results in the transfer of a significant part of the cellular polyamine pool to the nucleus (figure 6d) with particularly intense staining of RNA rich nucleoli. Additionally, SMLM of histone H3 indicates that in comparison to untreated cells (figure 6e), OND treatment (figure 6f) reduces the apparent density of chromatin associated histone H3 in the nucleus from 3813 (± 250) per µm^2^ to 842 (± 503) per µm^2^, whereas levels observed in the cytoplasm remain similar at 250 per µm^2^. Furthermore, the localization density achieved for total H3 is much lower than for DNA binding dyes, and is insufficient to discern OND induced chromatin compaction.

**Figure 6.**
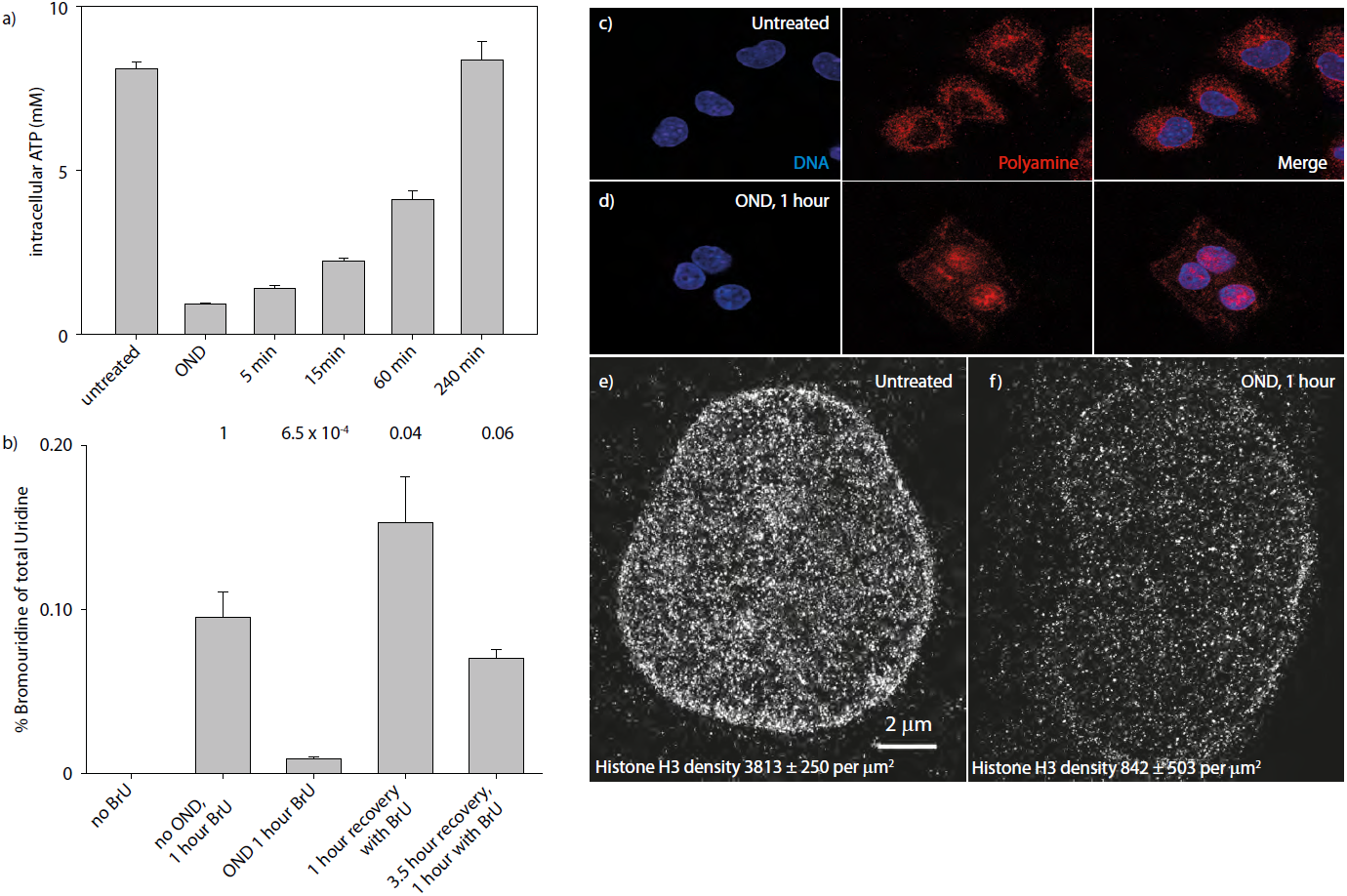
OND depletes intracellular ATP levels, inhibits transcription, induces relocation of the cellular polyamine pool to the nucleus and reduces the staining density of histone H3 with antibody. The intracellular concentration of ATP in untreated, OND exposed and in recovering cells was determined using a luciferase dependent assay (a). Global rates of transcription, determined by the incorporation of bromouridine into RNA, in untreated cells, cells under OND and cells recovering from OND are presented in (b). HL-1 cells, either untreated or subject to one hour of OND, were fixed, permeabilized and stained either with anti-polyamine antibody (c and d) or with anti-total H3 antibody (e and f) and counterstained with the fluorescent DNA binding dye Hoechst-33342. Cells were then examined using confocal microscopy. The content of immunostained histone H3 was evaluated by SMLM in untreated (e) and in OND treated (f) HL-1 cells.

### Fluorescent Recovery After Photobleaching (FRAP) indicates that core histones are not displaced from chromatin upon OND induced compaction, and that OND decreases the mobility of the linker histone H1

We wished to discriminate between possible explanations underlying the approximately 80% reduction in histone H3 staining upon OND treatment, as determined by SMLM. Potentially, this observation could arise through compaction restricting the accessibility of antibody to chromatin and / or through the direct loss of core histones from chromatin. By inference, histone loss from chromatin would liberate a highly mobile pool of histones, in contrast to their limited mobility when present in chromatin. We therefore used FRAP on live cells to estimate the mobility of Histone H2B labelled with mCherry in untreated HeLa cells and HeLa cells in an ischaemic environment. We selected H2B, which along with H2A, and in contrast to H3 and H4, exhibits significant exchange [46]. Consequently, FRAP analysis of H2B-mCherry is an appropriate marker for estimating OND induced displacement of core histones. As shown in figure 7a, HeLa cells undergo chromatin compaction when subject to one hour of OND, and significantly, H2B-mCherry retains a structured nuclear distribution, suggestive of chromatin compaction, indicating that a widespread release of core nucleosomes from chromatin does not occur upon OND treatment. FRAP measurements (figure 7b) of the mobility of H2B-mCherry confirm that OND does not increase the mobility of this core histone.

**Figure 7.**
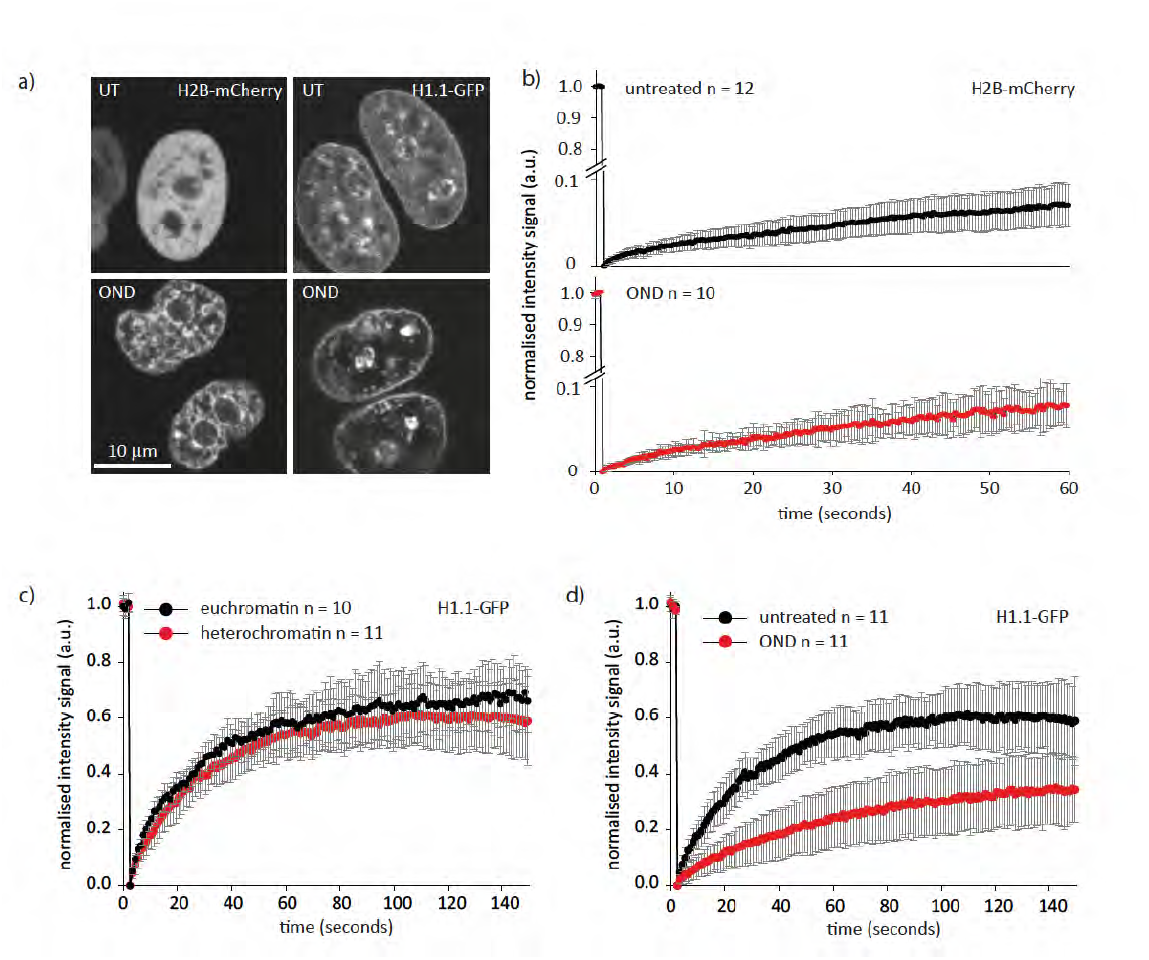
OND does not induce core histone displacement from chromatin but does decrease the mobility of the linker histone H1. We firstly demonstrated thatheLa cells stably transfected with either histone H2B-mCherry or with histone H1.1-GFP respond to one hour of OND by undergoing chromatin compaction. Comparison of untreated cells (top panels, figure 7a) with cells exposed to one hour of OND (bottom panels, figure 7a) by confocal microscopy clearly indicate that chromatin of HeLa cells compacts upon OND treatment. We then evaluated the mobility of the core histone H2B using FRAP on untreated (figure 7b, upper) and on OND treated (figure 7b, lower) cells. The recovery after photobleaching was extremely slow for both conditions, indicating that OND does not induce displacement of H2B from chromatin. We then evaluated the mobility of the linker histone H1 in untreated and in OND treated HeLa cells. As previously reported [47, 48], histone H1 is mobile, and is somewhat less mobile in heterochromatin than in euchromatin (figure 7c). As shown in figure 7d, OND induced chromatin compaction dramatically reduces the mobility of histone H1, indicating that the extent of chromatin compaction in OND is considerably higher than that between euchromatin and heterochromatin.

We then evaluated the mobility of the linker Histone H1.1 in HeLa cells, which maintains higher order chromatin structure through binding to extranucleosomal DNA. H1 exchanges continuously, with a residence time of a few minutes, even within heterochromatin [47, 48]. We firstly confirmed these observations in untreated HeLa cells, demonstrating that the mobility of Histone H1.1-GFP is higher in euchromatin as compared to heterochromatin (figure 7c). In agreement with the extensive compaction of chromatin induced by OND treatment for one hour, the mobility of Histone H1.1-GFP is significantly reduced upon treatment (figure 7d), demonstrating that (a) displacement of Histone H1.1 does not occur and that (b) OND induces chromatin compaction to an extent that restricts the exchange of Histone H1.1, and that the extent of this compaction exceeds the extent of the difference between euchromatin and heterochromatin. In conclusion, OND does not induce displacement of core histones but does reduce the mobility of the linker histone H1.1. This suggests that the decrease in density of immunostained H3 in OND results from compaction of a substantial fraction of chromatin to an extent that excludes penetrance by antibodies.

### OND induced chromatin compaction can be estimated by cytometry, provokes histone deacetylation and decreases the internal structure of cells

We further explored OND induced chromatin compaction using cytometric analysis of histone H3, and of post-translationally modified histone H3 variants. We reasoned that antibodies would stain compacted chromatin to a lesser extent than chromatin in untreated cells, thereby facilitating a semi-quantitative evaluation of the extent of OND induced histone chromatin compaction. Furthermore, ischaemia results in a general decrease of histone H3 [49-53] and H4 [54-56] acetylation levels. We therefore anticipated that OND should provoke a general reduction in antibody staining against histone marks, due to compaction restricting antibody access and moreover, that this effect should be especially pronounced for acetylated histone marks. In accordance with these considerations, OND induces a considerable reduction in staining of total histone H3, pan acetylated H3, H3K9ac, H3K14ac, H3K27ac, H3K4me3, and to a lesser extent, of H3K9me3 and of H3K27me3 (figure 8a). Accessible histone marks, such as acetylated H3 variants or trimethylated H3K4, are affected by OND to a greater extent as compared to either total H3 or, in particular, to histones present in compacted chromatin, such as H3K9me3 and H3K27me3. The kinetics of histone acetylation and methylation of lysine 14 of histone H3 is shown in figure 8b. One hour of OND induces a dramatic loss of H3K14 acetylation, which rapidly recovers upon restitution of oxygen and nutrients. H3K14 trimethylation exhibits little change through the experimental time-course. Similar results are obtained by confocal evaluation of cells stained with anti-H3K9ac (supplementary figure S4) and with anti-H3K14ac (supplementary figure S5); OND induces a profound loss of histone acetylation that recovers several minutes after release from OND. A useful feature of cytometric analysis is the detection of side-scattered blue light, which is proportional to the granularity or internal complexity of the cell. Side-scatter (SSC) is a measurement of mostly refracted and reflected light that occurs at any interface within the cell where there is a change in refractive index [57]. We anticipated that OND induced compaction of chromatin should result in a change in detected SSC, providing an independent methodology reporting the effect of OND upon chromatin. Importantly in the context of this analysis, OND does not induce a significant change in nuclear volume (supplemental results). As shown in figure 8c, OND induces a reduction in SSC that recovers on restoration of normoxia and nutrients. In keeping with our previous observations, side scatter measurements are significantly higher four-hours post-recovery as compared to untreated cells.

**Figure 8.**
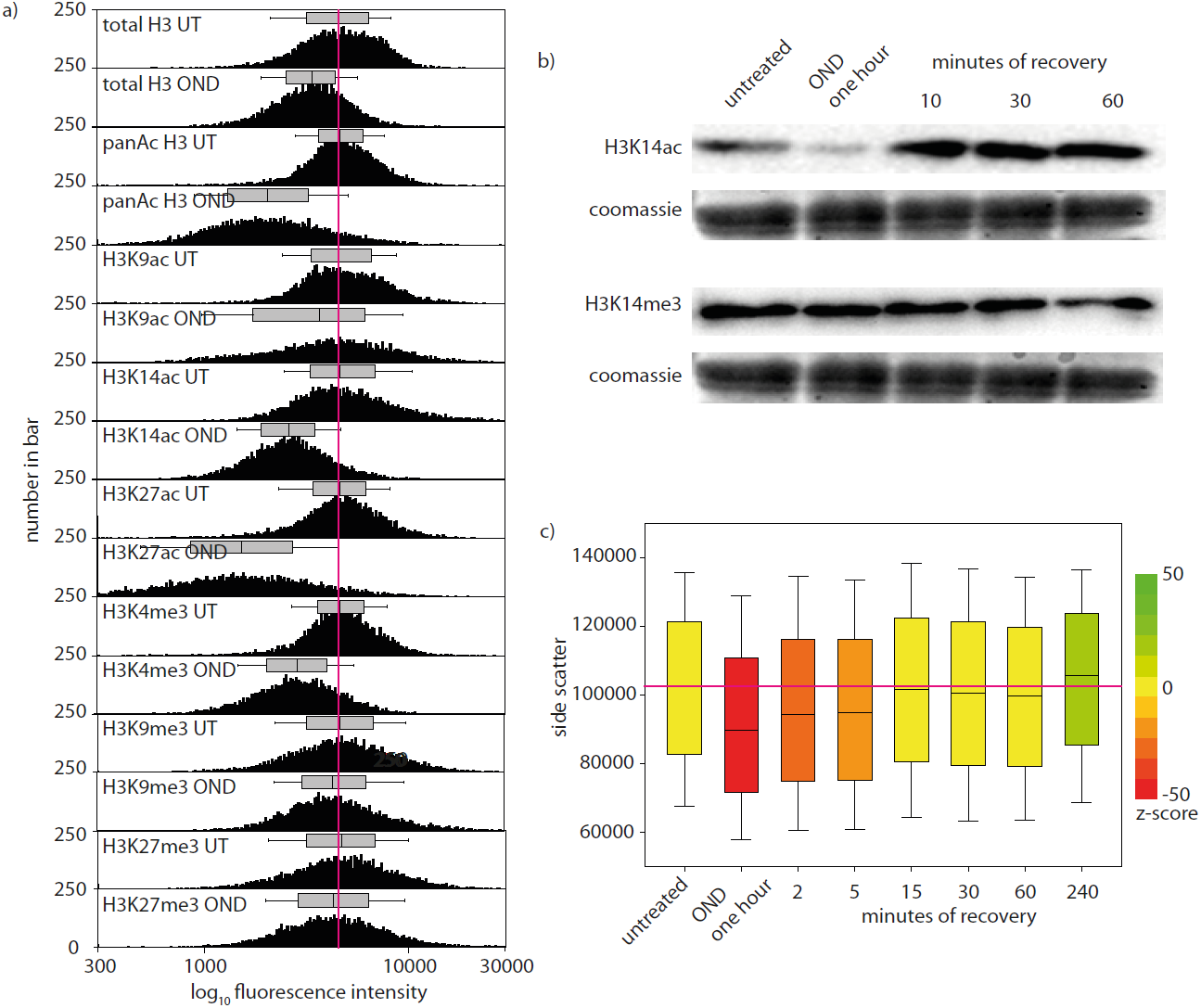
OND reduces access to chromatin by anti-histone antibodies, induces deacetylation of histones and a reduction in cellular granularity. HL-1 cells, either untreated or subject to one hour of OND, were trypsinised to produce a monodisperse suspension, fixed, permeabilized, washed and immunostained with anti-histone H3 antibodies, as indicated. Cytometric analysis was performed on a minimum of 10^4^ cells. A comparison of the staining intensity of untreated and OND cells is presented in (a); each data pair is normalized to the median of the untreated total H3 with the median and range of the distribution shown above each histogram. Western blot analysis of total H3K14ac and of H3K14me3 are shown in (b) across the experimental timecourse. The distribution of side-scatter measurements, which are proportional to internal cellular granularity, is presented for 10^4^ cells as box-plots showing the median, 25% and75% intervals as boxes and the 5% and 95% intervals as whiskers. The median value for untreated cells is shown as a horizontal line through all box-plots. The z-score between the untreated cell population and each other experimental condition, determined by the Mann-Whitney ranked sum test, is indicated by the colour of the box.

## Discussion

Ischemia is a defining event in the prevalent causes of morbidity in humans, including stroke, myocardial infarction and cancer. We show, using single molecule localization of DNA binding dyes, that the nuclear architecture of immortal cardiomyocytes, under experimental conditions mimicking transient ischemia, undergoes dramatic and reversible compaction. While functionally reversible compaction changes under ATP depletion conditions have previously been observed [32, 58, 59] using conventional microscopy approaches, this report quantitatively describes the nanoscale condensation of chromatin in myocardiac cells at single molecule resolution and has sufficient spatial resolution to qualitatively describe the extent of chromatin compaction. These analyses comprehensively reveal the extent, mechanism and reversibility of OND induced chromatin compaction and have been confirmed by alternative analytical procedures.

The extent of compaction indicates that chromatin undergoes a phase-transition under OND; that is, chromatin changes from a structurally more open, ‘disordered’ state to a structurally more closed, ‘ordered’ state. This is in keeping with the partial exclusion of DNA binding dye from chromatin upon OND and is consistent with the random obstacle network model recently proposed by Baum *et al.*, [60]. OND induces a previously undescribed sub-nuclear configuration comprised of discrete, DNA dense, atoll-like structures interspersed between large chromatin sparse voids. Moreover, OND provokes an extensive depletion of ATP and a relocation of the intracellular polyamine pool from the cytoplasm to the nucleus. Mechanistically, chromatin compaction is consistent with OND induced polyamine compaction and potentially, of divalent cation compaction, of chromatin. This process directly links cellular energy status with chromatin architecture. Upon cessation of ischemic-like conditions, the nuclear structure of cardiomyocytes undergoes relaxation, within a period of tens of minutes. Moreover, chromatin adopts a more open configuration, compared to untreated cells, several hours after release from OND. This effect may be of consequence in influencing the epigenetic reprogramming of cells.

In the presence of multivalent cations, high molecular weight DNA undergoes a dramatic condensation to a compact, usually highly ordered toroidal structure, with experimental evidence showing that DNA condensation occurs when about 90% of its charge is neutralized by counterions [61]. ATP exists in the cell predominantly as a complex with Mg^2+^. Consequently, OND mediated reduction in intracellular ATP concentrations increase intracellular availability of Mg^2+^ [62] and may promote chromatin compaction through divalent cation mediated charge shielding of phosphate groups in DNA. Experimentally increasing the osmolarity of culture medium [30] or increasing the exposure of detergent permeabilized cells to divalent cations, but not monovalent cations, provokes chromatin compaction as evaluated by either confocal microscopy or by FLIM-FRET [32]. Similarly, as the ATP-Mg^2+^ complex sequesters intracellular polyamines, principally spermine and spermidine [63], a reduction in ATP levels will result in the intracellular polyamine pool transferring to chromatin, by mass-action, thereby further enhancing condensation [32] and displacement of histones. In keeping with these proposed effects, transient ATP depletion by inhibition of oxidative phosphorylation with azide in SW13 and in HeLa cells induces in increase in the volume of the interchromosomal compartment, as observed by confocal microscopy [58].

Chromatin primarily consists of DNA wrapped around the core histone complex [64]. Additionally, OND results in a profound loss of active histone marks, particularly acetylation and H3K4 trimethylation. This raises the issue of how the histone code, particularly for active genes, is reinstated during recovery from an ischemic environment and may provide new insight into the phenomenon of ischemic preconditioning, where pretreatment of an organ with short periods of ischemia has a protective effect on subsequent ischemic insult [65]. The degree of chromatin relaxation upon recovery from OND exceeds that of untreated cells, indicating that chromatin may adopt a more transcriptionally permissive configuration as compared to either untreated or OND treated cells. This could arise in consequence of intracellular ATP levels exceeding the divalent and polycation pool of the cell upon recovery from OND, such that chelation of the intracellular polyamine and divalent cation pool by ATP exceeds the amount present under continuous normoxic and nutrient-rich conditions.

There is an intimate connection between chromatin architecture and the functional output of chromatin. Advanced technologies are revolutionizing understanding of chromosome organization, and are progressing understanding of spatial organization on transcription, replication and repair [66]. Ischaemic conditions provoke an extreme level of chromatin compaction, as witnessed, for example, by the extensive development of chromatin free areas within the nucleus and in the restriction of linker histone H1 activity mobility that far exceeds that within heterochromatin. However, the methodologies we describe, if utilised in conjunction with specific labelled genomic regions, could be developed to probe local DNA configurations. The defining characteristic of the SMLM we have performed is the labelling density that can be achieved using DNA binding dyes. This allows resolution of chromatin nanostructure at a scale appropriate to inform on regulatory events.

The extent and reversibility of chromatin compaction induced by OND suggests that the impact of ischemia could be constrained by targeting biochemical events that are required for chromatin condensation. In this light, pan-inhibition of histone deacetylase (HDAC) activity is efficacious in animal models of cerebral ischemia [67] and specific knockdown of HDAC3 or HDAC6 promotes survival of cortical neurons in an *in vitro* model of ischemia employing oxygen and glucose deprivation [68]. Increased HDAC activity was reported in a mouse model of cardiac ischemia and inhibition of HDACs by Trichostatin A treatment significantly reduced the infarct size [69]. Moreover, OND induced compaction of chromatin may explain the observed increase of histones in serum that occurs in catastrophic ischemic events. Alternative strategies could be to chelate intracellular divalent cations or limit the production of polyamines, for example through inhibition of ornithine decarboxylase activity. Intriguingly, these is extensive pre-clinical evidence that this strategy is of benefit in restricting the growth of solid tumours [70] and inhibiting ornithine decarboxylase protects *Drosophila* against hypoxia induced reduction in lifespan [71]. In summary, ischemic conditions induce a rapid compaction of chromatin, which is associated with a general inhibition of transcription [6]. Correspondingly, nuclear architecture senses and responds to environmental conditions, through structural rearrangements. Defining and understanding these effects offers a diverse range of tractable targets for therapeutic intervention in human disease.

## Conclusions

- Experimental ischemia induces compaction of chromatin
- Chromatin compaction is reversible upon restoration of normoxia and nutrients
- Experimental ischemia lowers ATP levels and reorganizes polyamines into the nucleus
- Upon recovery from compaction, chromatin acquires a more open and transcriptionally active configuration

## Methods

### Cells and cell culture

HL-1 cells are an immortal mouse cardiomyocyte cell line, derived from a murine atrial tumour, which retain the morphology and gene expression profile of adult cardiomyocytes and the ability to contract [72]. HL-1 cells were cultured in gelatin/fibronectin-coated dishes in Claycomb medium (Sigma) supplemented with 2mM glutamine (Gibco), 0.1mM norepinephrine (Sigma-Aldrich), 10% FBS (Sigma-Aldrich) in 5% CO_2_, 37°C and 95% humidity. Cells were passaged every 3 days as described [72]. For microscopic analysis, HL-1 cells were grown on coated glass cover slips (Assistant, 20×20 mm) in 6-well plates to a density of 50%. All other experiments were performed on confluent cells.

### Oxygen Nutrient Deprivation (OND)

HL-1 cells were washed twice with PBS (Gibco) and placed in a hypoxia chamber (Whitley Hypoxystation H35) with 1% O_2_, 5% CO_2_, 94% N_2_ at 37°C and 70-85% humidity. An ischemic environment was simulated by incubating cells in 115mM NaCl, 12mM KCl, 1.2 mM MgCl_2_, 2mM CaCl_2_, 25mM HEPES and 5mM deoxyglucose; this solution was pre-equilibrated to 1% O_2_ prior to use. Cells were incubated for 1 hour under these conditions, after which they were washed with PBS then returned to Claycomb media under normoxic conditions. Experimental evaluation was usually performed on untreated cells, cells subject to one hour of OND and on recovering cells at 5, 15, 60 and 240 minutes following OND. Recovery was designed to mimic reperfusion after an ischemic event. Non-recovered OND treated cells were harvested and fixed in a hypoxic atmosphere. All buffers used for the preparation of such samples were equilibrated to 1% O_2_ in advance. Untreated cells were kept under normal culture conditions until fixation.

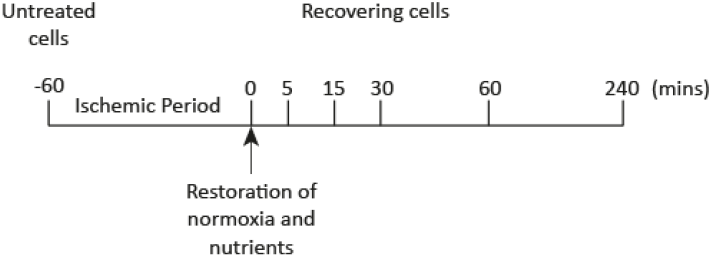

Schematic representation of the time-course employed to evaluate the effect of OND, and subsequent recovery, on HL-1 cells.

### Immunofluorescence analysis of histones, Lamin B1 and DNA by confocal microscopy

Cells grown on coated cover slips in 6-well plates were treated with 2 ml of OND buffer or with Claycomb medium as indicated. Cells were then washed twice with PBS, fixed in 1 ml ice cold methanol for 10 min, washed with PBS and permeabilized with 1 ml PBS containing 0.3% Triton X-100 (Sigma-Aldrich) and 0.3% Tween-20 (Sigma-Aldrich) for 10 minutes at room temperature. Blocking was performed with 1 ml blocking buffer (5% BSA and 0.1% Tween-20 in PBS) for 1 hour at room temperature. For the antibody labelling, cells were incubated with anti-H3 (Abcam, 1 µg/ml), anti-H3K14ac (Cell Signaling, 1:500), anti-Lamin B1 (Abcam, 1 µg/ml) or anti-polyamines (Abcam, 10µg/ml) overnight at 4 °C in 500 µl blocking buffer. After incubation with primary antibody, cells were washed three times with 1 ml wash buffer (PBS containing 0.1% Tween-20) and incubated with AlexaFluor 488 conjugated secondary antibody (Invitrogen, 2 µg/ml)) for 1 hour in 1 ml wash buffer, followed by three washes with wash buffer. DNA was stained with Hoechst 33342 (0.5 µg/ml) for 20 minutes at room temperature and washed three times with 1 ml PBS. Cells were then embedded in 10 µl of glycerol. For analysis, a Leica SP5 II confocal system (Leica Microsystems GmbH) with a 63x oil immersion NA1.4 objective lens was used, and 1024x1024 images were acquired using a pinhole size of 1.0 Airy units, 60-100 nm pixel pitch.

### Fluorescence Recovery After Photobleaching (FRAP)

FRAP experiments utilised HeLa cells stably transfected with mCherry-H2B or with GFP-H1.1. Live cell experiments were performed either in OND-buffer or in RPMI 1640 without phenol red (Life Technologies) containing 10% foetal bovine serum (Gibco). OND samples were tightly sealed using Picodent Twinsil two component glue (Wipperfuerth, Germany) within a hypoxia chamber prior to FRAP analysis. The following parameters were used for acquisition of FRAP data:

|  | **H2B-mCherry** | **H1.1-GFP** |
| --- | --- | --- |
| **FRAP image acquisition** | 256 x 256 pixel,<br>effective pixel size 96.5 nm,<br>excitation: 6% power of 561 nm,<br>emission: 602nm - 711 nm | 256 x 256 pixel,<br>effective pixel size 96.5 nm,<br>excitation: 13% power of 488 nm,<br>emission: 500nm - 591nm |
| **bleaching protocol** | one 256 x 256 pixel scan in 2 x 2 μm, 80% 561 nm laser | 100 ms point bleach, 80% 488 nm laser |
| **time/frame** | 0,328 sec | 1 sec |
| **FRAP read-out in:** | 2 x 2 μm square ROI | 1.5 μm in diameter circle ROI |

Data were processed as described in Trembecka-Lucas *et al.* [73] with some modifications. Eleven individual FRAP measurements were performed for each data set. Each FRAP acquisition was aligned using the StackReg ImageJ plugin, to prevent from movement of bleached area during fluorescence recovery [74]. Fluorescence recovery after photobleaching was then analysed within a respective region of interest using ImageJ and corrected on bleaching throughout the experiment by acquisition of bleaching curves for either H1.1-GFP or H2B-mCherry in independent measurements in both buffers, to accomodate possible differences in bleaching rates.

### Flow cytometry analysis of histone marks

Cells were grown to confluency in 10 cm diameter culture dishes, then were subject to ischemia/reperfusion protocol as previously indicated, using 10 ml of OND solution or Claycomb medium. Cells were then washed twice with 10 ml PBS, trypsinized with 1 ml 0.25% Trypsin (Gibco) and vigorously resuspended in 5 ml of PBS containing soybean trypsin inhibitor. The cells were centrifuged at 250 xg for 5 minutes, washed once in 10 ml PBS and the cell pellet was resuspended in 1 ml ice cold methanol for 10 minutes to fix the cells. Cells were again centrifuged, and then permeabilized for 10 minutes in PBS containing 0.3% Triton X-100 and 0.3% Tween-20. After a further centrifugation, the cell pellet was resuspended in 500 µl PBS and the cell density was estimated using an automated cell counter (BioRad). One million cells were resuspended in 300 µl FACS buffer (PBS with 0.1% Tween-20, 1% BSA) containing anti-H3 (Abcam, 3 µg/ml), anti-H3K14ac (Cell Signaling 1:300), anti H3K9ac (Cell Signaling, 1:300), anti-H3K27ac (Abcam, 3 µg/ml), anti-pan-acetylated H3 (Merck Millipore, 3 µg/ml), anti-H3K4me3 (Merck Millipore, 3 µg/ml) and anti-H3K27me3 (Active Motif, 3 µg/ml) and incubated for 1 hour. Cells were washed three times with 1 ml FACS buffer and incubated with 1 µg/ml AlexaFluor 488 conjugated secondary antibody (Invitrogen) for 45 minutes. After a further three washes with FACS buffer, cells were resuspended in 300 µl PBS and the fluorescent intensity of the cell population was analysed using a BD LSRFortessa cytometer (BD Biosciences). Cells were gated so that only events from discrete single cells were counted, with 10^4^ events recorded per experimental condition.

### ATP determination

Cells were grown to confluency in 3.5 cm diameter culture dishes and oxygen and nutrition deprivation (OND) applied to selected samples as described previously, using 2 ml of ischemic salt solution as required. After OND, cells were washed twice with PBS, and ATP extracted with boiling water as described by Yang et al. [75]. ATP was determined using a bioluminescence assay utilizing recombinant firefly luciferase and its substrate D-luciferin (ATP Determination Kit, Invitrogen).

### Bromouridine labeling of nascent RNA

Newly synthesized RNA was pulse labeled by incubating confluent 10 cm culture dishes with 2 mM Bromouridine (BrU) (Sigma Aldrich) for 1 hour, either under normal culture conditions, under OND or for 1 hour of recovery from OND. To estimate background incorporations, cells were analysed without exposure to BrU. All experimental conditions were performed in triplicates. Following BrU pulse labeling, RNA was extracted using Trizol (Ambion) following the manufactures instructions and following digestion to nucleosides with nuclease P1 (Roche), U snake venom phosphodiesterase (Worthington) and alkaline phosphatase (Fermentas) as described by Kellner *et al.* [76]. RNA nucleosides were subject to LC/MS analysis. Separation was performed on an Agilent 1290 UHPLC system equipped with ReproSil 100 C18 column (3 µm, 4.6 × 150 mm, Jasco GmbH) maintained at 30°C. Identification and quantification of nucleosides was performed on an Agilent 6490 triple quadruple mass spectrometer.

### Determination of nuclear volume

Cells were cultured in 6-well plates on coverslips and subject to one hour of OND as described. Cells were then fixed for 10 minutes on ice with methanol, permeabilized for 10 minutes in PBS containing 0,3% Triton and stained with Hoechst 33342 (2µg/ml, Sigma). Samples were embedded in glycerol and nuclear volume was calculated by reconstruction of nuclear z-stacks following acquisition on a Leica SP5 II confocal system (Leica Microsystems GmbH) using a step size of 0.21 µm. A 1.4 NA 63× oil objective was used. The Imaris software package (Bitplane) was used to calculate nuclear volume using the following parameters: surface grain size: 0.170 µm, threshold absolute intensity: 14.6702, distance to image border xy: 0.429µm, volume: above 200 µm. Statistical analysis was performed using the Mann-Whitney Rank Sum test.

### Western Blot

Histones were extracted from 10 cm dishes of confluent HL-1 cells after OND and OND plus 10, 30, or 60 minutes of recovery. The cells were washed with PBS supplemented with 5mM sodium butyrate to prevent deacetylation. The cells were collected in 800 µl of Triton extraction buffer containing PBS with 0,5% Triton X-100, 2mM phenylmethylsulfonyl fluoride (PMSF) supplemented with 5mM sodium butyrate and protease inhibitor. Cytoplasma lysis was performed for 10 minutes on ice followed by a 10 minutes centrifugation step at 2000 rpm at 4 °C. The resulting nuclei pellet was resuspended in 100 µl of 0.2 N hydrochloric acid and histone extraction was performed over night at 4 °C and rotation. After a 2000 rpm centrifugation step the supernatant was collected and protein concentration was determined by Bradford assay. Five microgram of histones were diluted in laemmli loading buffer and boiled for 5 minutes. The samples were run on a 12.5 % SDS gel and subsequently blotted on nitrocellulose membranes for 1 hour at 100 V at 4 °C. Ponceau staining was used as a loading and transfer control. The membranes were blocked in 5 % bovine serum albumin in TBST buffer for 1 hour, incubation with primary antibody (H3K14ac, abcam, 1:5000, H3K14me3, Signalway Antibody, 1:5000) was done overnight at 4 °C. After three TBST washes the secondary antibody conjugated to horse radish peroxidase was incubated for 45 minutes and washed off three times. The blots were developed with 1 ml ECL reagent (Invitrogen) per blot and pictures were taken under a ChemiDoc (Biorad).

### DNAseI digestion of chromatin

Time-course *in situ* digestion assays, to determine the relative resistance of chromatin in untreated and in OND treated cells to DNAseI, were performed using Leica SP5 confocal microscope (Leica, Wetzlar, Germany) at 37^o^C. HL-1 mouse were seeded on IBIDI 8-well chamber, subjected to 1 hour of OND, or not, then fixed for 15 min with 4% PFA, followed by permeabilisation using 0.3% TX-100 in PBS. Cells were stained with 320 µl of 5 µM DRAQ5 (Life Technologies) for 30 min then subsequently washed twice with PBS. PBS was replaced with 150 µl 1× DNase buffer (NEB) and placed on to the microscope stage. 150 µl of 10 U/ml DNase I in DNase buffer was then (5 U/ml final concentration) and time-lapse measurement of DRAQ5 fluorescence was initiated. Images were taken every 4 minutes with autofocus stabilization, and were acquired with following settings: 7% 633 nm laser excitation, 643 - 749 nm emission range, 512 × 512 resolution (voxel size 246 × 246 × 481.5 nm^3^), 2AU pinhole, 600 Hz scanner speed.

### Localization Microscopy

#### Sample preparation for Single Molecule Localization Microscopy (SMLM)

HL-1 cells on coverslips were cultured, fixed and permeablilized as described. Samples were washed twice with PBS then incubated for 40 min with 1 µM Vybrant dye cycle Violet (Life technologies), followed by a further two washings with PBS. For SMLM imaging of YOYO-1, cells on coverslips were permeabilized, incubated with 0.5 U/ml RNase A and 20 U/ml RNase T1 (Ambion, USA) for 1 h at 37^o^C, then stained with 0.02 nM YOYO-1 in 1 ml PBS for 30 min. Cells were then washed twice with PBS and embedded in 20 µl PBS containing 40 µg/ml glucose oxidase, 0.5 mg/ml catalase and 10% (w/v) glucose. SMLM was performed after 300 minutes, once the majority of dye dissociates from the DNA to the interior of the nucleus (Supplementary Figure S7). For SMLM imaging of AlexaFluor 647 stained samples, an imaging buffer of 40 µg/ml glucose oxidase, 0.5 mg/ml catalase, 10% (w/v) glucose and 50 mM (for Lamin B1 staining) or 100 mM (for histone imaging) mercaptoethylamine (MEA) in 60% (v/v) glycerol and 40% (v/v) PBS was used. For SMLM imaging of Vybrant dye cycle Violet an imaging buffer consisting of 40 µg/ml glucose oxidase, 0.5 mg/ml catalase, 10% (w/v) glucose in 80% glycerol and 20% PBS was used. For two color imaging of DNA and AlexaFluor 647, the imaging buffer was further enriched with 3 mM MEA (a concentration that facilitates blinking of the cyanine dye AlexaFluor 647 without affecting blinking of Vybrant dye cycle Violet (Żurek-Biesiada *et al*. 2015, submitted). 20 µl of the appropriate buffer was placed on a glass slide after which the coverslip with fixed cells was placed upside-down on the droplet. The coverslip was bonded to the slide with biologically inert, impervious, dental paste (Picodent Twinsil) prior to SMLM imaging.

#### SMLM measurements

The SMLM configuration has been described previously [36]. Briefly, the custom built microscope is equipped with a single objective lens (Leica Microsystems, 1.4 NA, 63× oil immersion with a refractive index of 1.518) and a 12 bit, air-cooled CCD camera (PCO, Sensicam QE, effective pixel-size in the sample region corresponds to 102 nm). For fluorescence discrimination, emission filters used for Vybrant dye cycle Violet, YOYO-1 and AlexaFluor 488 were bandpass filter 525/50 nm (Chroma) and for AlexaFluor 647 long pass 655 nm (Chroma). Widefield acquisitions are performed with homogeneous illumination of the whole field of view, as achieved by 6.7 fold expansion of the laser beam. SMLM images are acquired upon illumination with a collimated laser beam covering an area of approximately ∼25 µm diameter in the imaging plane (full width at half maximum of gaussian profile). For single color imaging of AlexaFluor 647, a 647 nm laser (LuxX diode laser, Omicron, Germany) was used at 60 mW (measured in a sample plane), and 25,000 frames with an exposure time of 25 ms were acquired for each SMLM experiment. Stained DNA (Vybrant dye cycle Violet, YOYO-1) in HL-1 cells was excited using a 491 nm laser (Calypso 05 series, Cobolt, Sweden). For Vybrant dye cycle Violet 30,000 frames with 50 ms exposure time were acquired at 70 mW laser power (sample plane), and for YOYO-1 30,000 frames with an exposure time of 50 ms were acquired at 30 mW laser power (sample plane). In the dual color experiments, imaging of AlexaFluor 647 (9,000 frames with 25 ms exposure time, 647 nm excitation, 60 mW in the sample plane) was performed prior to imaging of DNA stained with Vybrant dye cycle Violet. For imaging of AlexaFluor 488 23,000 frames with camera exposure time of 25 ms were acquired at 70 mW 491 excitation.

#### Data analysis and post-processing

SMLM data analysis was performed using fastSPDM, a custom software package written in Matlab (Mathworks) (Gruell et al., 2011) to extract single molecule fluorophore positions from raw tiff data stacks. First, an initial background image was calculated by averaging over eight frames. Noise in the background was calculated assuming a Poisson noise model, i.e. the standard deviation of the noise is given by STD = (background)^1/2^. Looping through each of the acquired frames, the background was subtracted from the raw data, yielding a difference image. Initial estimation of the positions of single fluorophore signals were detected with pixel accuracy in the difference images (after smoothing with a 3 × 3 mean filter) based on a Threshold Factor (TF), defined as follows: Only signals with a peak intensity I0 higher or equal to (TF−1)*STD were considered. Subpixel accuracy positions were extracted from a 7×7 ROI around the initial estimation of each signal by calculating the center of intensity. Overlapping signals detected as radially increasing intensity as we move away from the center pixel of the ROI were clipped, yielding a refinement of the positions. If clipping results in > 30% loss of accumulated signal within the ROI, then the signal is discarded. From the remaining signals, a list of localizations was generated containing information on x,y-position, photon count, and localization precision σ_loc_ [77] for each signal, where localization precision of a point emitter is defined as follows:

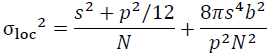

where *N* corresponds to number of photons, *s* is a standard deviation of Gaussian point-spread function, *p* is a pixel size, *b* is a background noise. The difference image was further used to adjust the background image for the next frame as follows: Values larger than STD were clipped from the diff image, and the result was scaled by a factor of 1/8 and added to the previous background image. This new estimate of the background image was made positive by clipping values less than 0.

For analysis of the SMLM data, TF values of 3 (DNA stained with Vybrant dye cycle Violet or YOYO-1) or 3.5 (AlexaFluor 647) respectively were used. The initial list of localizations as described above was altered by joining single molecule signals occurring in consecutive frames (search radius = 2.5 <σ_loc_>). No further filtering of the list of localizations e.g. based on localization precision, was performed. Such filtering of the list of localized signals in PALM/STORM related approaches based on the width of the detected signal is often necessary because even weak signals are easily picked up if the background intensity is close to zero. Assuming a Poisson noise model for the detected photons, then also the noise in the detected background is close to zero. In our approach, single molecule signals of the DNA dyes are detected on top of a relatively high background (average of 150 counts, i.e. 300 photons). In this case, the x-y-broadening of the detected fluorescence signals as we move away from the focal plane is sufficient to reject these out-of-focus signals, as they become hidden in the noise level.

Prior to final reconstruction, the drift in the images occurring during acquisition was determined from the list of localizations using correlation of reconstructions of up to 100 subsets. The drift determined in this way amounts to approximately 150 nm/h, and was used to correct the list of localizations. The shift between drift-corrected subsets of data usually did not exceed a standard deviation of

*σ*_*drift*_= 10 nm.

For visualization/reconstruction of the SMLM data, Gaussian blurring was used: All single fluorophore positions were blurred with their respective localization precision σ_*loc*_ (AlexaFluor 647) or with the average localization precision <σ_loc_> (DNA/SMLM data) to generate a SMLM reconstruction. All reconstructed images were generated with a pixel size corresponding to 5 nm.

The single molecule photon count of DNA dyes has an average of about 1500 photons, i.e. much less than the brightest dSTORM/SMLM fluorophores. The resolution *σ*_*total*_ in the DNA SMLM data was estimated from the photon count dependent localization precision using the formula

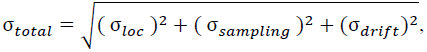

Where *σ*_*loc*_ is the average localization precision [77], 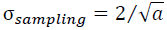 where a is the average density of localized single molecules of fluorophores (typically between 4,000 - 6,000 molecules / µm^2^). Our data generated a theoretical value of average 2D resolution, σ_total_, of 39 nm, resulting in an effective structural resolution of about 90 nm, assuming a normal distribution of the measurement error. In addition, by means of Fourier Ring Correlation [78], we calculated the lateral resolution in our studies to be approximately 100 nm. The single molecule photon count of DNA dyes has an average of about 1500 photons, i.e. much less than the brightest dSTORM/SMLM fluorophores. Single molecule signals of the DNA dyes, as observed in an optical section through a cell nucleus are detected on top of a relatively high background. In such a case, the SMLM lateral resolution will be determined mostly by the noise in the background, and not reach as good values as reported manifold for SMLM [36].

When performing dual-color SMLM imaging, the chromatic shift within cell samples was determined from SMLM experiments of immunostained microtubule standards (double labelling with primary and mixture of Atto 488-/AlexaFluor 647-coupled secondary antibodies), using the same emission filter set as in the DNA experiments. The shift in the two emission channels of the microtubule data was extracted and then used to correct single fluorophore coordinates from two different detection channels in two-color DNA experiments.

For 2D histogram analysis of stained DNA by SMLM, a square grid of variable width was placed over the cell, and a Matlab function was used to count localized molecules within each square (bin) of the grid. In order to characterize the spatial change in chromatin upon OND, the grid width (bin size) was varied between 10 nm and 500 nm. Subsequently, a histogram of localized single fluorophores per bin was plotted, and the median and quartile values of localizations were determined. This allows us to quantitatively compare the difference in density distribution for control, OND and recovering cells. Nine or more cells were evaluated for each experimental condition. Bins containing less than two localized molecules were discarded from the analysis (DNA-free area in particular from outside the nucleus).

Analysis of DNA binding dye-free area in 2D SMLM images of Vybrant dye cycle Violet stained cells was performed using ImageJ (http://imagej.nih.gov/ij/) as follows: Reconstructed 8-bit images were subject to histogram stretching in order to cover the entire spectrum of pixel values. Then, a constant threshold for all images analyzed was applied, after which the 8-bit image was converted to a binary image. The area covered by chromatin was calculated by counting the number of pixels in this binary image. Similarly, the total area of the nucleus in the imaging plane was obtained after the chromatin devoid “holes” were filled (ImageJ function “fill holes”). Then the chromatin free area was calculated by subtracting the chromatin occupied area from total nuclear area.

### Structured illumination (SIM) of YOYO-1 stained DNA

For SIM imaging HL-1 cells were treated as described above for SMLM imaging. Cells were stained with YOYO-1 and immediately embedded in Vectashield H-1000 (refractive index 1.45, VectorLabs). We used 488 nm excitation and camera integration times between 200 - 300 ms. A sinusoidal illumination pattern was generated in the focal plane by interference of laser light, resulting in a grid pattern of 280 nm spacing. Three different orientations of the grid with three different phases for each orientation were used, resulting in 9 acquired images per 2D slice. The microscope setup and reconstruction software has been described [79, 80].

## List of abbreviations

ac: acetyl
AMPK: Adenosine MonoPhosphate Kinase
ATP: Adenosine TriPhosphate
BALM: Binding Activated Localization Microscopy
CD: Chromatin Domain
CDC: Chromatin Domain Cluster
CoA: Coenzyme A
DNA: DeoxyriboNucleic Acid
EdU: 5-ethynyl-2’-deoxyuridine
FLIM-FRET: Fluorescence Lifetime Imaging Microscopy - Förster Resonance Energy Transfer
FRAP: Fluorescent Recovery After Photobleaching
H: Histone
HIF: Hypoxia Inducible Factor
IC: Interchromosomal Compartment
K: lysine
me: methyl
OND: Oxygen and Nutrient Deprivation
PR: Perichromatin Compartment
SAM: S-adenyl Methionine
SSC: SideSCatter.

## Author Contributions

C.C. and G.R. conceived this study and designed the experimental strategy. I.K., A.S., C.H., I.C., M.M., D.M. and K.F. performed experiments. I.K., A.S., K.P., F.S., M.M., U.B. and G.R. analysed data. U.B. prepared the movie. U.B., C.C. and G.R. supervised this project. G.R. wrote the paper and coordinated the supplemental information. All authors discussed the results and contributed to the manuscript. No author has any conflict of interest with respect to publication of this work.

## Acknowledgements

We wish to thank Prof. William Claycomb, LSU Health Sciences Center, New Orleans, for his generous gift of the HL-1 cell-line, the Cytometry Core Facility and the Microscopy Core Facility at the Institute of Molecular Biology, Mainz, for their assistance in generating data, and Profs Hiroshi Kimura, Prof. Jurek Dobrucke and Dr. Miroslaw Zarebski for the H1.1-GFP HeLa cell-line. This work was supported by the Boehringer Ingelheim Foundation. We wish to thank Dr. Phillip Wenzel, and Profs. Thomas Cremer, Frank Gannon and Kurt Kremer for useful discussions.

## Additional material

We have prepared a separate supplemental file containing a supplementary results section, reporting in detail experimental results briefly referred to in the main manuscript, a supplementary note describing artefacts in localization microscopy of chromatin, a supplementary movie illustrating how our SMLM was performed and given the interdisciplinary nature of our work, a glossary.

Supplementary Movie M1. Generating a small molecule localization nanograph. The steps taken to generate a SMLM image are illustrated for a single cell nucleus that has been exposed to one hour of oxygen and nutrient deprevation then fixed, permeabilized and its DNA labeled with Vybrant DyeCycle Violet. The left-hand window shows the raw data obtained every 50 ms, along with an indication of the laser power used to illuminate the sample. The upper-centre panel indicates the background level of fluorescence, which is subtracted from the raw-data profile to generate the signal above background frame (lower centre). The right-hand window contains the processed localizations, which are integrated to generate a localization map describing chromatin structures at sub-diffractional resolution. Low laser power was used for the first 100 frames to establish background fluorescence, thereafter high laser power was used to bleach most Vybrant DyeCycle Violet molecules. Continued application of high laser power induces a low number of optically isolated molecules of fluorophore to a bright/ON state, as a result of photoswitching. These are filtered to meet criteria of brightness and chastity and their localization accuracy determined, based on the number of photons emitted by the fluorophore. Signals above background are used from frame 200 onwards to contribute to the SMLM map. The next 600 frames (42 seconds) are shown contributing to the SMLM reconstruction. Thereafter, the final reconstructed image, consisting of 25,000 frames (imaging for 28 minutes and 44 seconds) is shown.

